## Supplemental Information for "Non-Invasive Single-Cell Morphometry in Living Bacterial Biofilms"

**Figure S1**

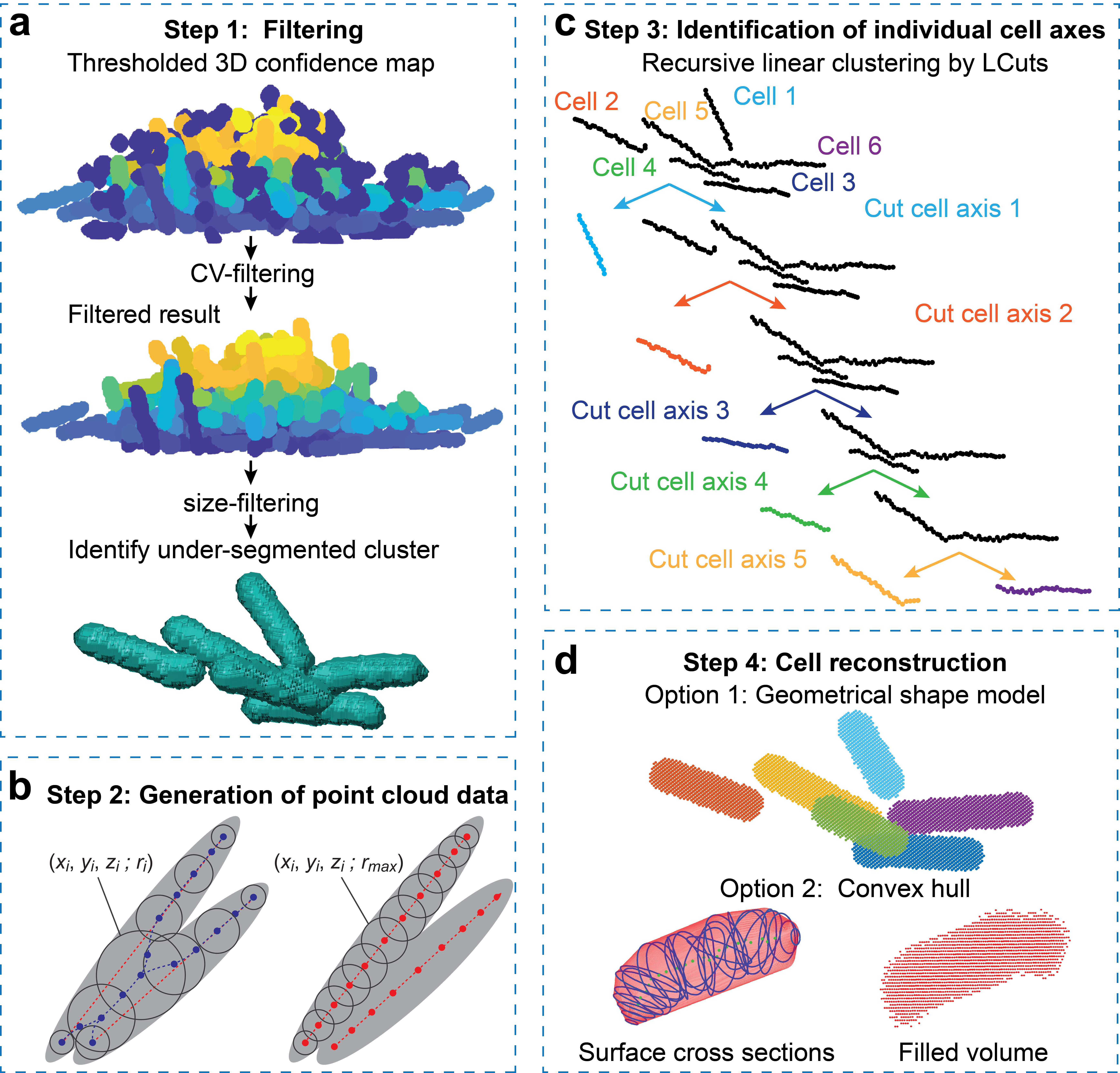

**Figure S1.** **Post-processing of CNN-produced confidence maps using a refined *LCuts* processing pipeline.** (**a**) False positive objects are detected and removed by CV- and size- filtering. Under-segmented clusters that are larger than single cells are selected for further splitting. **(b)** Illustration of modified medial axis (red dashed lines) extraction to generate point cloud data from fused clusters of rod-shaped cells using the method of inscribed spheres. When cells are touching, the traditional medial axis extraction process fails to align with the actual cell central axis (left). To overcome this drawback, we limited the size of the inscribed spheres based on prior knowledge of bacterial cell diameters (right). **(c)** The set of inscribed sphere centers are then treated as a fully-connected, undirected graph in 3D with two node features: location and direction (see text and **Figure S12** for details). The graph (blue nodes) is then iteratively cut into smaller graphs (red nodes) until the stopping criteria are reached (see text for details). **(d)** Post-processed graphs represented in different color denoting different cells. The 3D surface of individual cells can be determined using a geometrical cell shape model (e.g. a spherocylinder for rod shaped bacteria) or by calculating the convex hull around the inscribed spheres found in step 2.

**Figure S2**

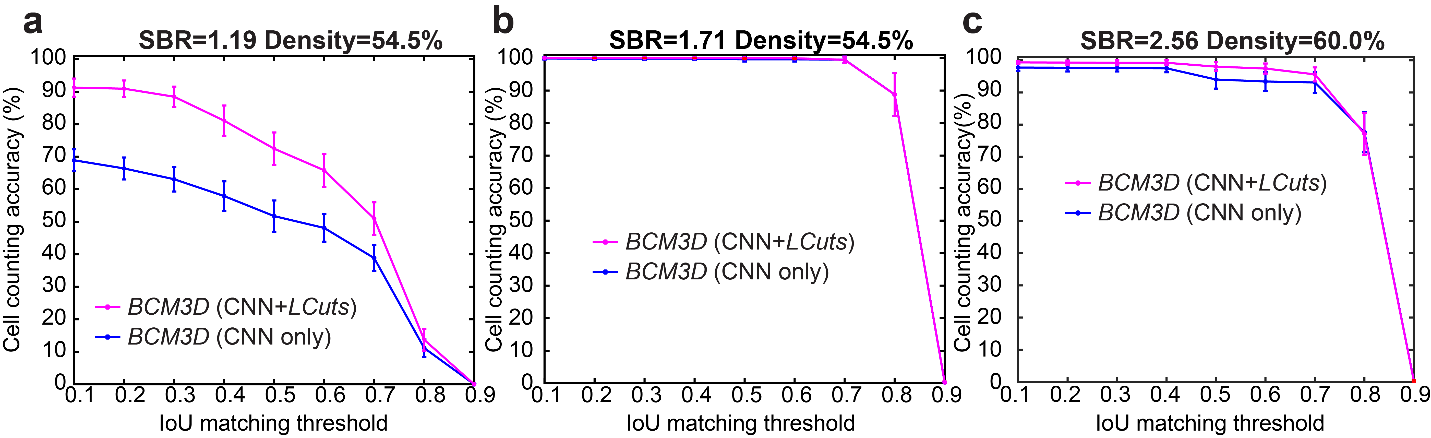

**Figure S2.** **Segmentation accuracies achieved for biofilm images with different SBR and cell densities.** (**a**), (**b**), and (**c**) represent results for datasets with SBR 1.19, Density 54.5%, SBR 1.71, Density 54.5% and SBR 2.56, Density 60.0%, respectively. Each curve is plotted by averaging 10 different datasets. Error bars represent ± one standard deviation.

**Figure S3**

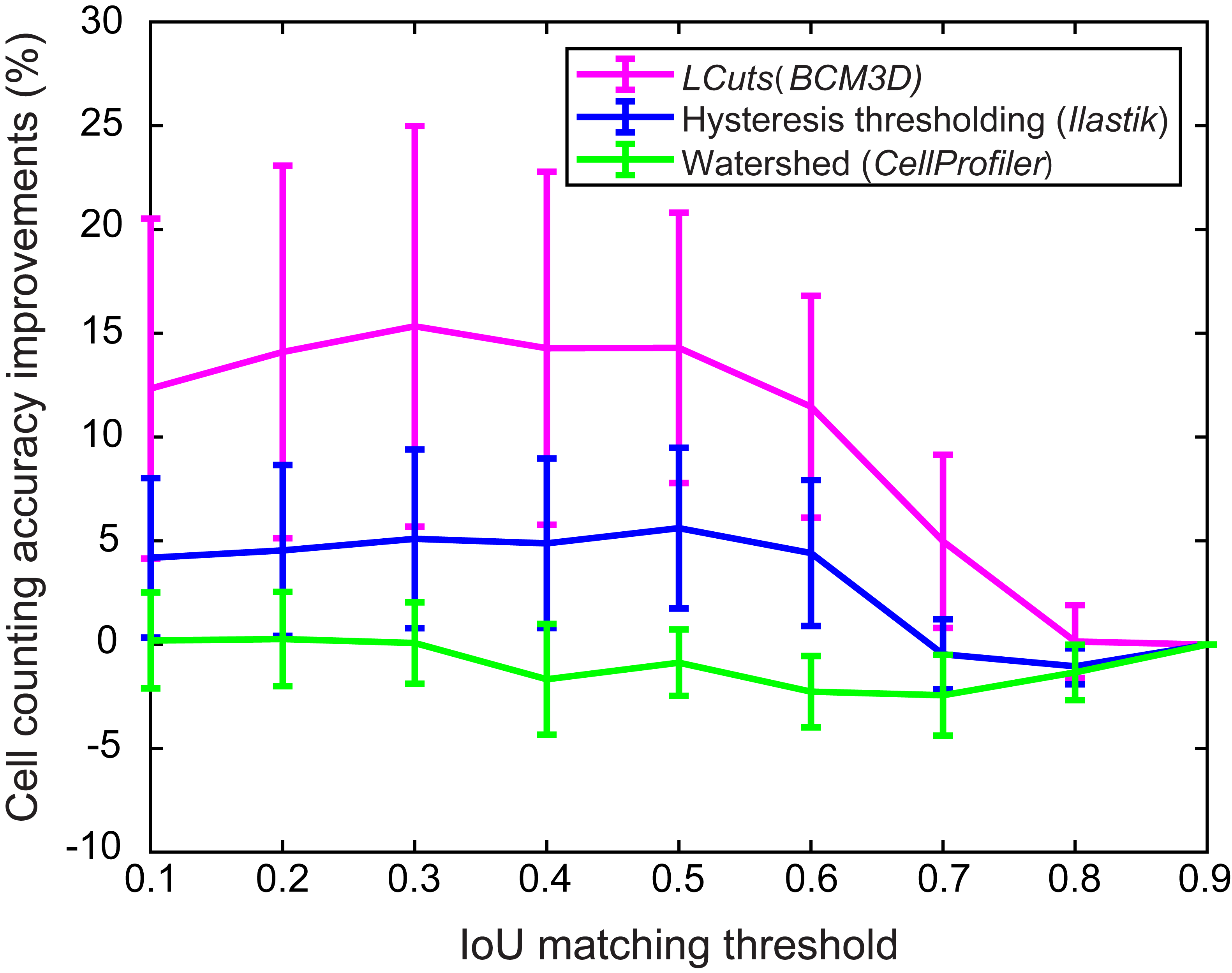

**Figure S3.** **Comparison of *LCuts* to commonly used image post-processing methods.** Shown is the cell counting accuracy averaged *N* = 20 randomly chosen, simulated datasets of low SBR and/or high cell density, for which post-processing is required. The hysteresis thresholding-based algorithm of *Ilastik*^1^ improves the cell counting accuracy by less than 6% on average for IoU matching thresholds less than 0.6. On the other hand, the watershed-based pipeline used by *CellProfiler*^2^ provides negligible improvements and even decreases the average cell counting accuracy in many cases. This decrease is primarily due to oversegmentation. Among the three methods tested, *LCuts* provides the highest improvement in cell counting accuracy (>12% on average for IoU matching thresholds less than 0.6). Data are presented as mean values ± one standard deviation indicated by error bars.

**Figure S4**

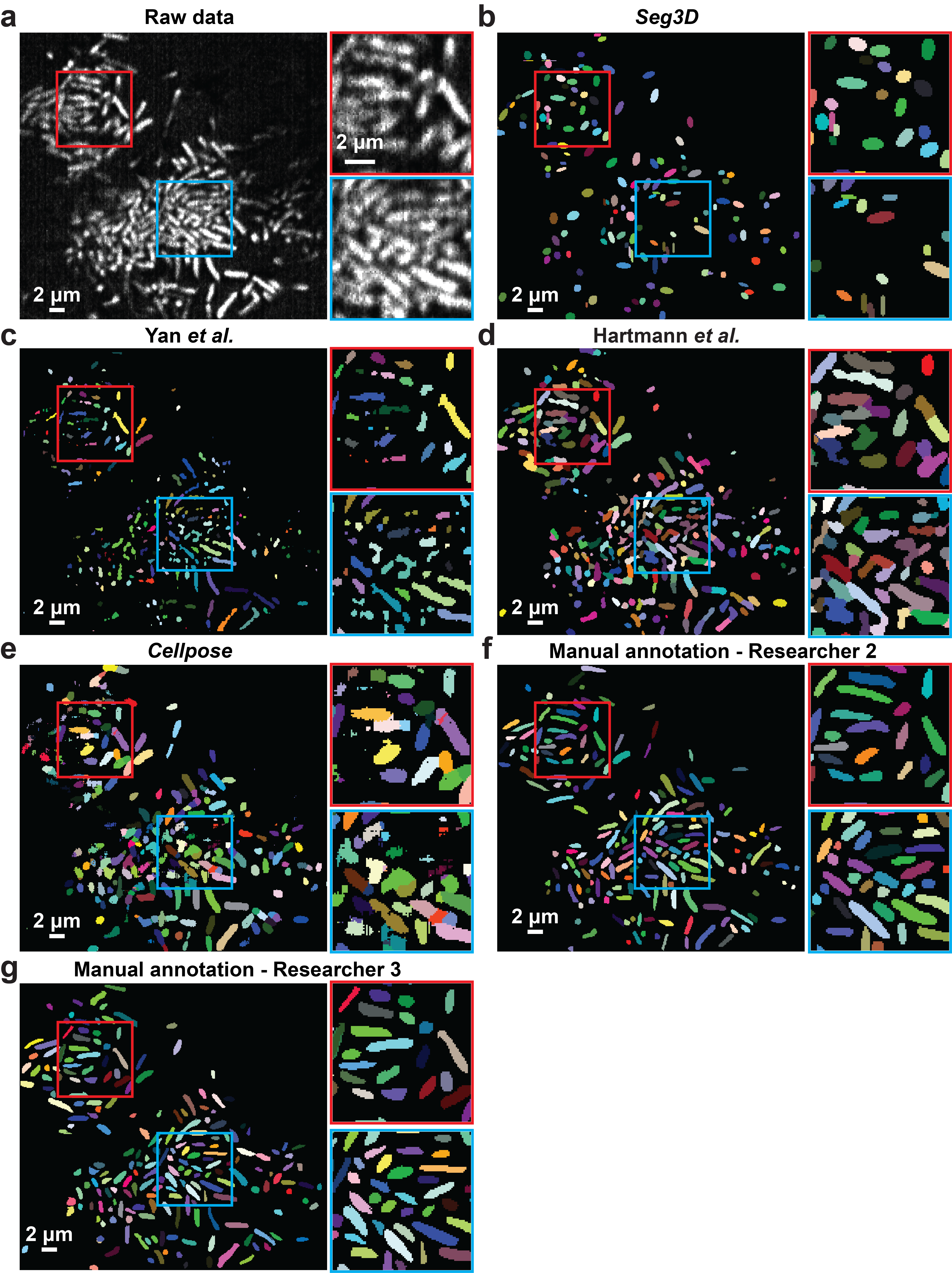

**Figure S4.** **Visual comparison of segmentation results achieved by previous segmentation approaches.** (**a**) *E. coli* biofilm image of GFP expressing cells 10 hours after inoculation (same image as in **Figure 4d**, reproduced here for comparison). (**b**) Segmentation result obtained using *Seg3D*. (**c**) Segmentation result obtained using the algorithm in Yan *et al*^3^. (**d**) Segmentation result obtained using Hartmann *et al.*^4^. (**e**) Segmentation result obtained using *Cellpose*^5^. Similar results were also obtained at the *t* = 300 and *t* = 360 time points.

**Figure S5**

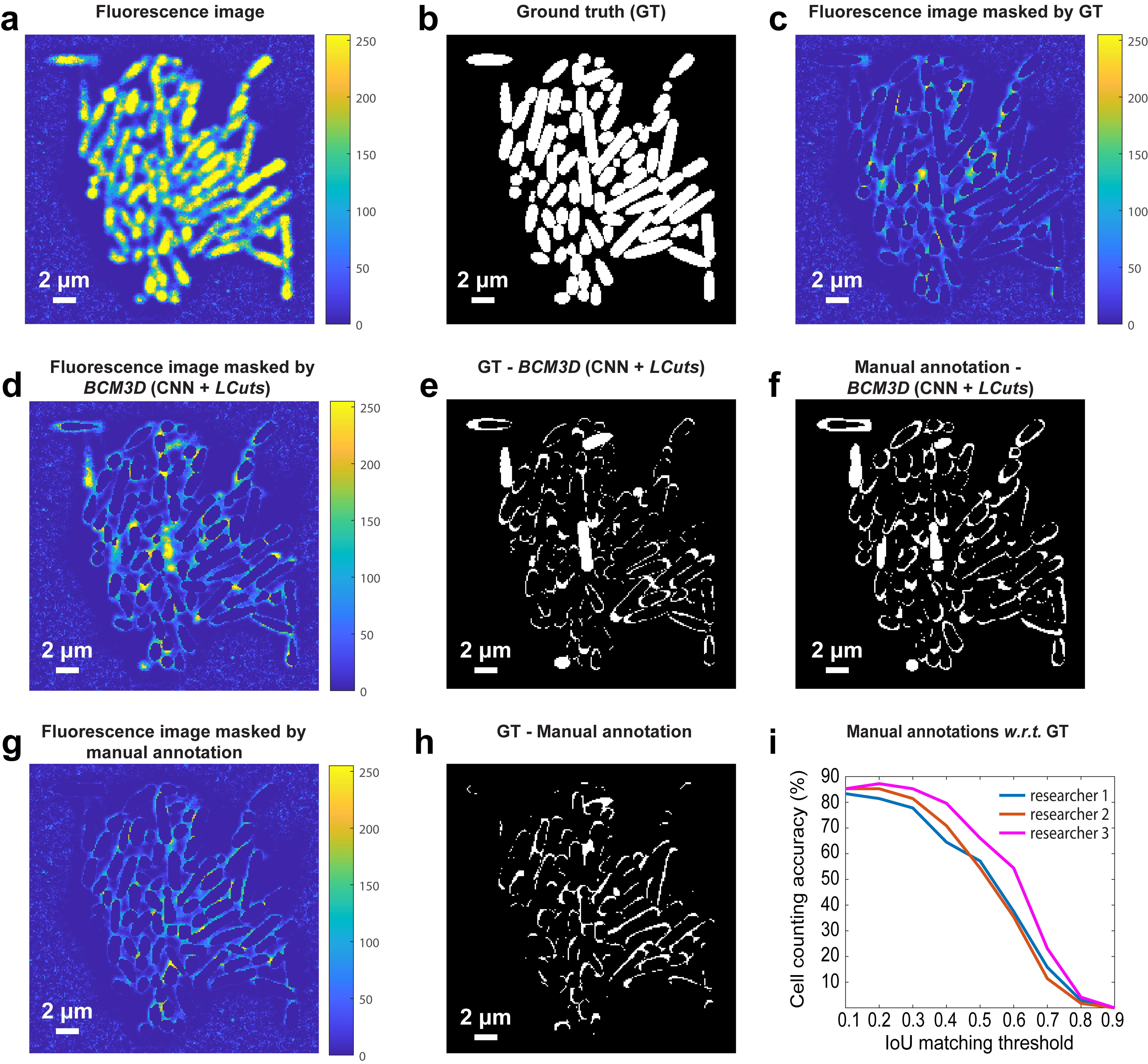

**Figure S5 Differences between manual annotation and ground truth. (a)** Fluorescence image slice of a 3D simulated biofilm and **(b)** the corresponding ground truth (GT). **(c)** The fluorescence image slice shown in (a) masked by its corresponding GT shown in (b). The fluorescence is not completely masked because of the diffraction-limited resolution of light microscopy. **(d)** Fluorescence image slice of the same simulated biofilm masked by the *BCM3D* segmentation result. **(e)** Absolute value of the difference image between the GT and the *BCM3D* segmentation result. White pixels indicate regions where the two masks do not agree. **(f)** Absolute value of the difference image between a manual annotated mask (from researcher 3) and the *BCM3D* segmentation result. **(g)** Fluorescence image slice of the same simulated biofilm masked by the manual annotation result. Researcher 3 chose to draw larger cell boundaries to mask more of the fluorescence intensity. **(h)** Absolute value of the difference image between the GT and the manually annotated mask. White pixels indicate regions where the two masks do not agree. *N* = 10 independent images with similar cell densities, but different cell arrangements were analyzed with similar results. **(I)** Segmentation accuracy achieved by manual annotation performed by three different researchers. Segmentation accuracy is parameterized in terms of cell counting accuracy (*y* axis) and IoU matching threshold (*x* axis, a measure of cell shape estimation accuracy. Curves approaching the upper right-hand corner indicate higher overall segmentation accuracy with respect to the ground truth. While IoU matching thresholds <0.3 yield good cell counting accuracies, the cell counting accuracy sharply decreases for IoU matching thresholds >0.3, because manually annotated cell shapes differ from the ground truth cell shapes.

**Figure S6**

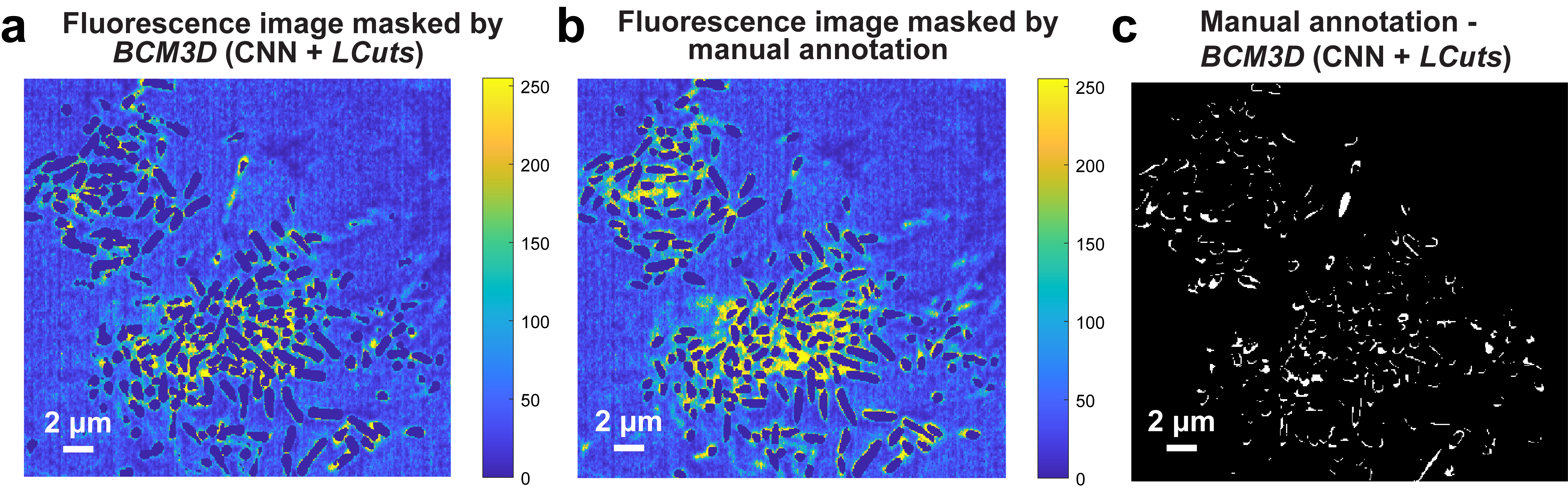
**Figure S6.** **Differences between manual annotation and *BCM3D* segmentation. (a)** Fluorescence image slice of the 3D *E. coli* biofilm shown in Figure 5d masked by the *BCM3D* segmentation result. **(b)** Fluorescence image slice of the 3D *E. coli* biofilm shown in Figure 5d masked by manual annotation. **(c)** Absolute value of the difference image between manual annotation and *BCM3D* segmentation. Similar results were also obtained at *N* = 3 different time points.

**Figure S7**

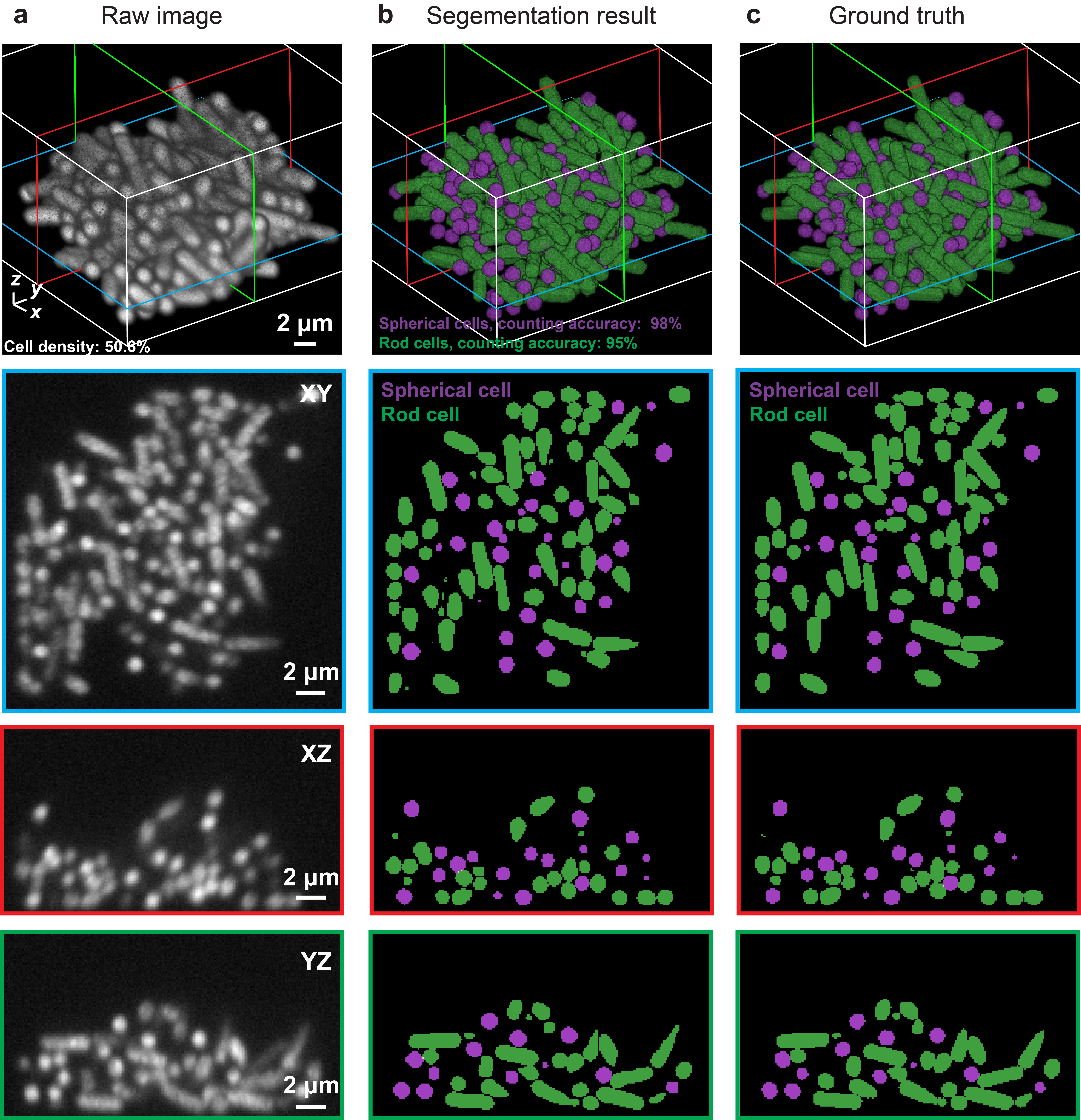

**Figure S7.** **Segmentation of mixed population biofilms containing spherical cells and rod-shaped cells.** (**a**) Simulated fluorescence image of a mixture of spherical cells and rod-shaped cells. The mixing ratio for this particular biofilm is 50:50, the cell density is 50.6%, and the SBR is 2.56. (**b**) Segmentation result obtained by *BCM3D* and (**c**) ground truth. Three orthogonal planes are shown below each 3D image. *N* = 10 independent images with similar cell densities, but different cell arrangements, where analyzed for each data point in **Figure 6**.

**Figure S8**

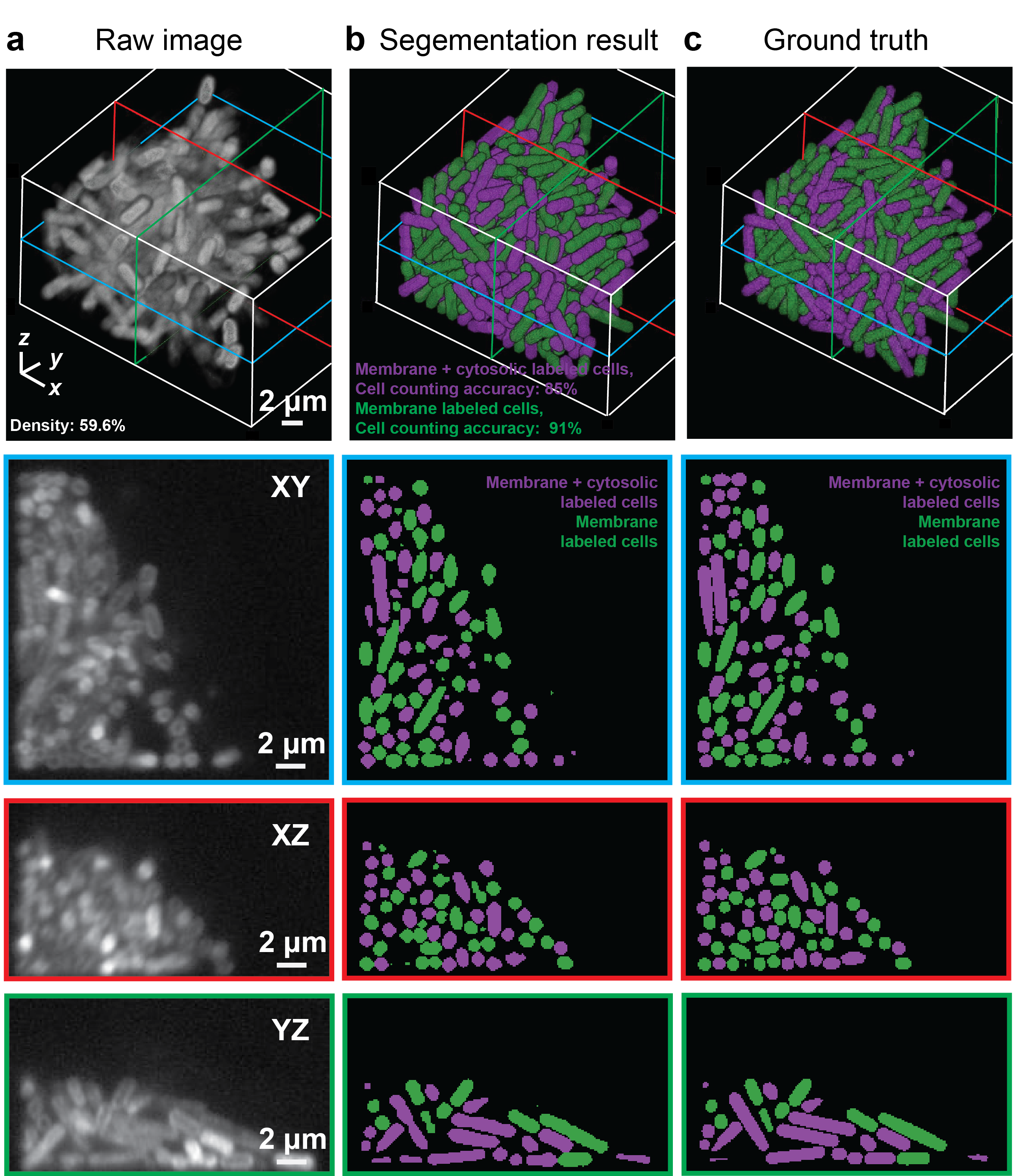

**Figure S8.** **Segmentation of simulated, mixed-population biofilms containing membrane-stained cells and membrane-stained cells that additionally express an intracellular fluorescent protein.** (**a**) Simulated fluorescence image of a mixture of membrane-stained cells and membrane-stained cells that additionally express an intracellular fluorescent protein. The mixing ratio for this particular biofilm is 50:50, the cell density is 59.6%, and the SBR is 2.56. (**b**) Segmentation result obtained by *BCM3D* and (**c**) ground truth. Three orthogonal planes are shown below each 3D image. *N* = 10 independent images with similar cell densities, but different cell arrangements, where analyzed for each data point in **Figure 6**.

**Figure S9**

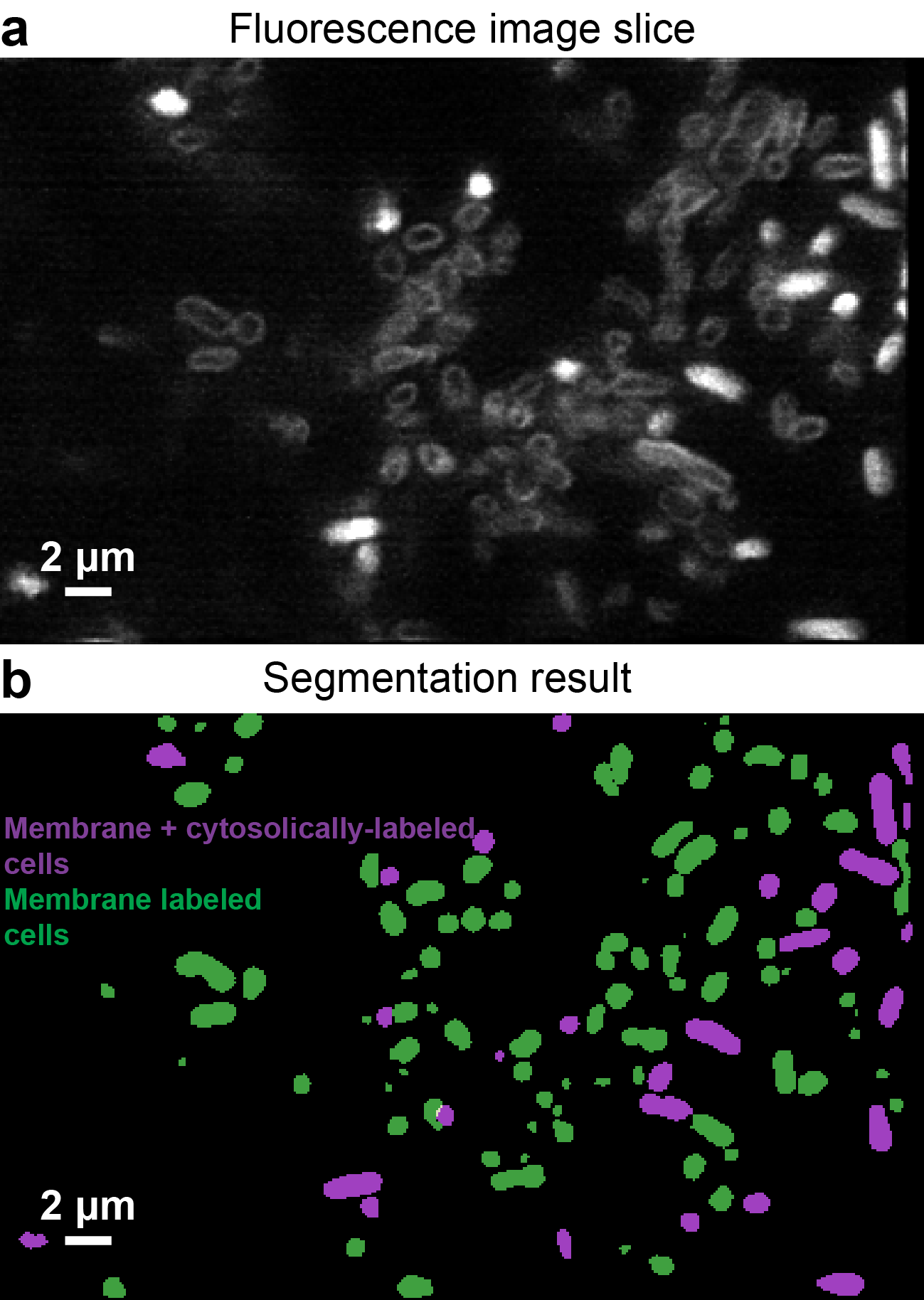

**Figure S9.** **Segmentation of experimental, mixed-population biofilms containing membrane-stained cells and membrane-stained cells that additionally express an intracellular fluorescent protein.** (**a**) Experimental 2D slice of a mixed *E. coli* population containing membrane-stained cells and membrane-stained cells that additionally express an intracellular fluorescent protein. The mixing ratio at the time of inoculation was 50:50. All cells were labeled by the FM4-64 membrane-intercalating dye. **(b)** *BCM3D* segmentation result corresponding to the image shown in (a). Membrane-stained cells are displayed in green, and cells that were both membrane-stained and cytosolically-labeled are displayed in magenta. Similar results were also obtained in *N* = 3 independent experiments.

**Figure S10**

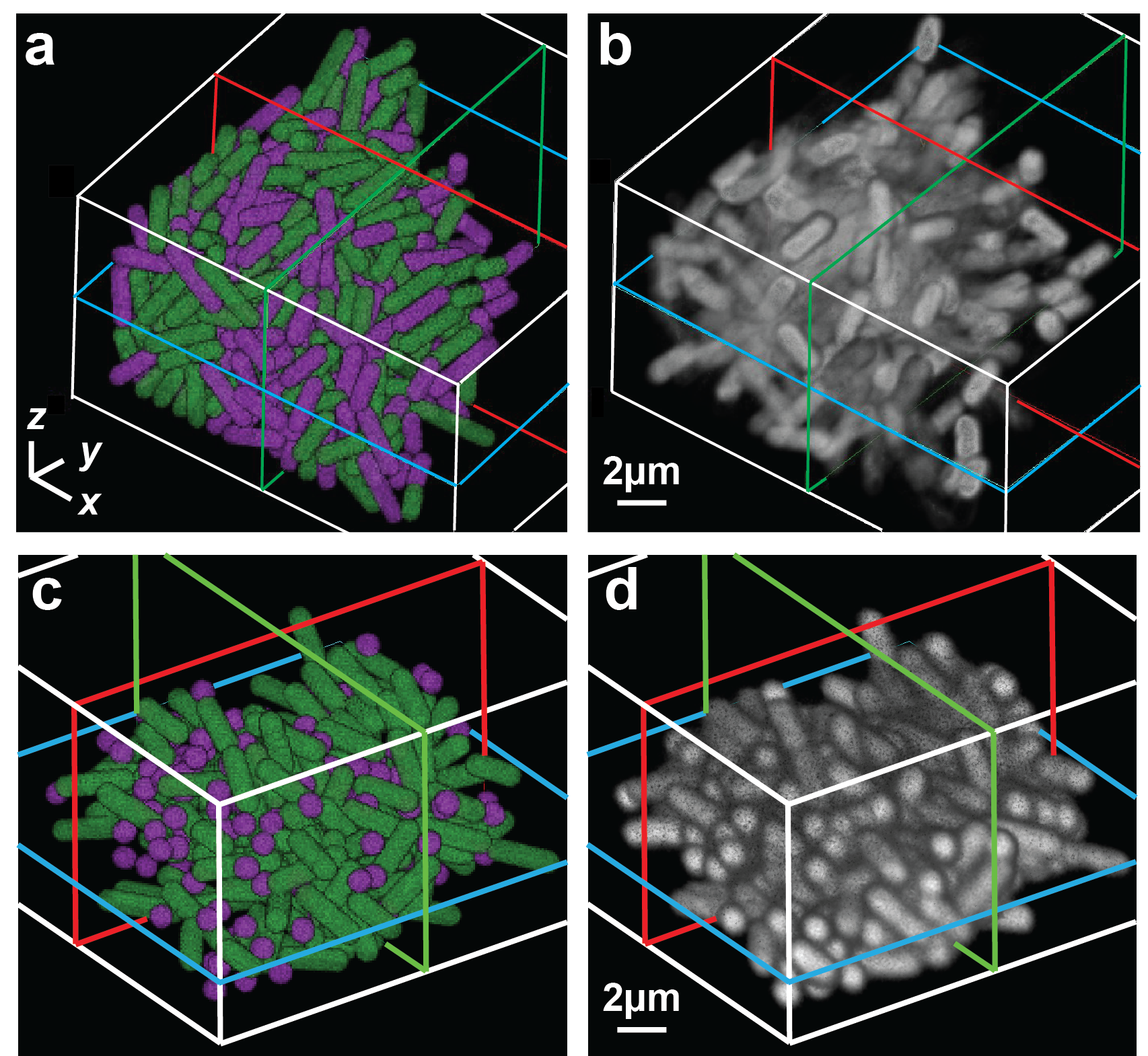

**Figure S10.** **Simulation of mixed labeling and mixed cell shape biofilms.** Cell arrangements (green indicates membrane labeled cells, magenta indicates membrane labeled cells that simultaneously express interior fluorescence protein). (**b**) Simulated fluorescence image based on the cell arrangements in (a) as displayed by the volume viewer plugin of Fiji^6^. *N* = 10 independent images with similar cell densities, but different cell arrangements, where analyzed for each data point in **Figure 6**. (**c**) Cell arrangements (green indicates rod-shaped cells, magenta indicates spherical shaped cells). (**d**) Simulated fluorescence image based on the cell arrangements in (c) as displayed by the volume viewer plugin of Fiji^6^. *N* = 10 independent images with similar cell densities, but different cell arrangements, where analyzed for each data point in **Figure 6**.

**Figure S11**

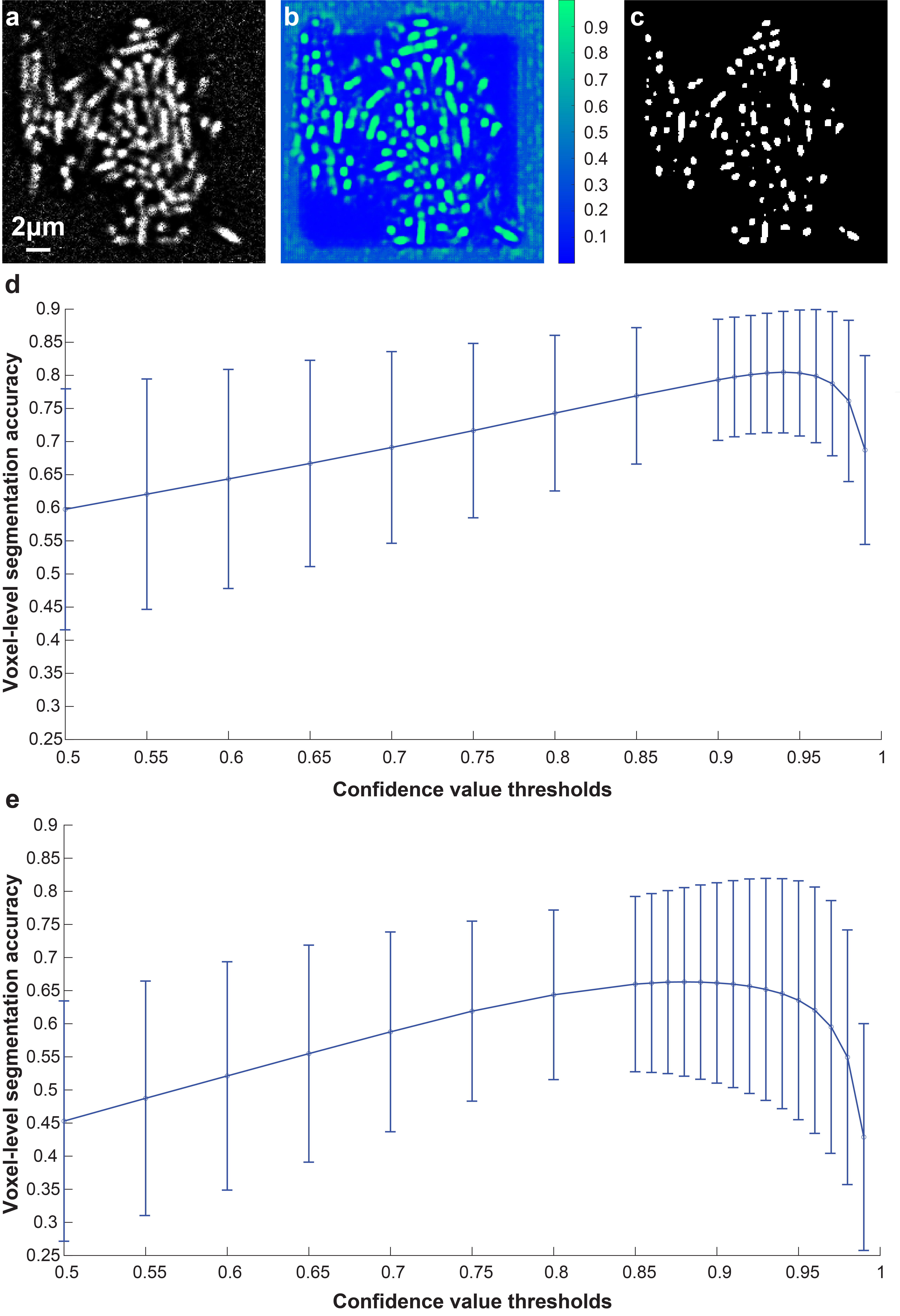

**Figure S11.** **Estimation of ‘cell interior’ confidence map threshold values.** (**a**) Deconvolved fluorescence image of the simulated biofilm in **Figure 1**. (**b**) ‘Cell interior’ confidence map. (**c**) Binary segmentation result (confidence threshold = 0.94). (**d** and **e**) Voxel-level segmentation accuracy (y axis) versus the confidence value thresholds (*x* axis) for cells labeled with cytosolic fluorophores (d) and cells labeled with membrane-localized fluorophores (e). Each curve is plotted by averaging *N* = 500 datasets, similar to those in panel (a), but with different cell densities and SBRs. Data are presented as mean values ± one standard deviation indicated by error bars.

**Figure S12**

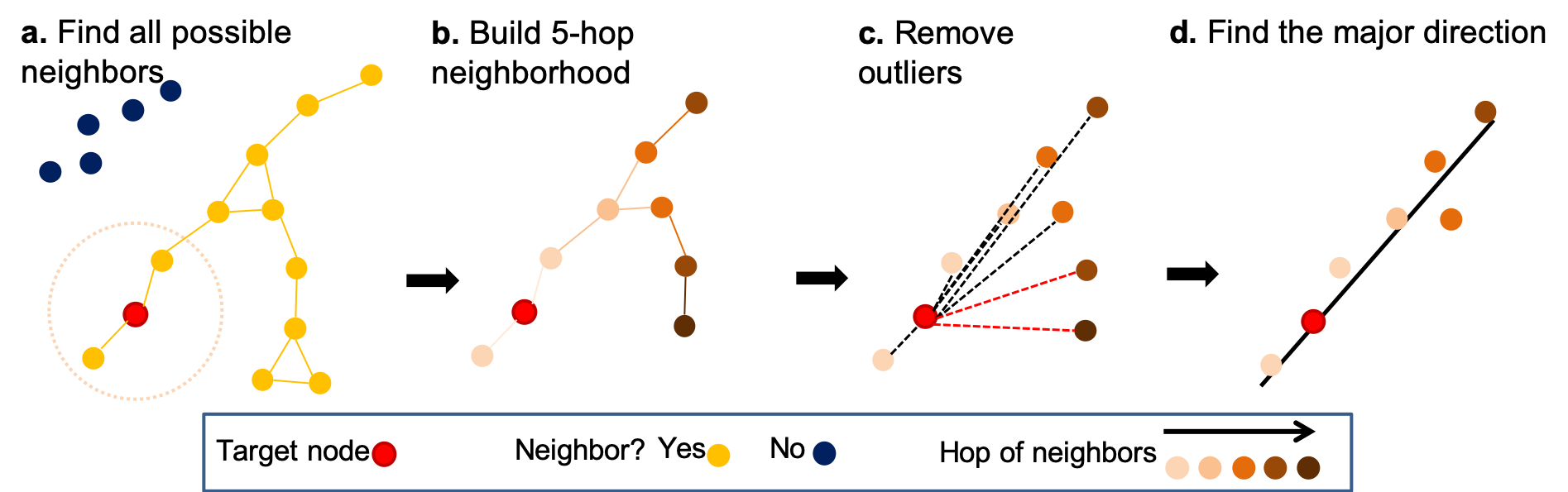

**Figure S12.** **Determination of node direction in an outlier-removed neighborhood**^7^. (**a**) A neighborhood of the target node (in red) is a sub-graph, where all adjacent nodes (in yellow) are connected via edges to the target node. Here, if the distance of two nodes is less than a chosen value (indicated by the dashed circle), these nodes are adjacent to each other. The blue dots are not part of the neighborhood. (**b**) A hop is defined as the number of edges that one has to traverse from one node to the other node in the graph. Here, the 5-hop neighborhood of the target node is shown. (**c**) The directional vectors are found from the target node to all the other nodes within in the 5-hop neighborhood (dashed lines). The nodes are classified as outliers if they have large relative angles compared to all the other directional vectors (red dashed lines). (**d**) Finally, the direction feature of the current node is evaluated as the major direction of the outlier removed neighborhood using principle component analysis.

**Figure S13**

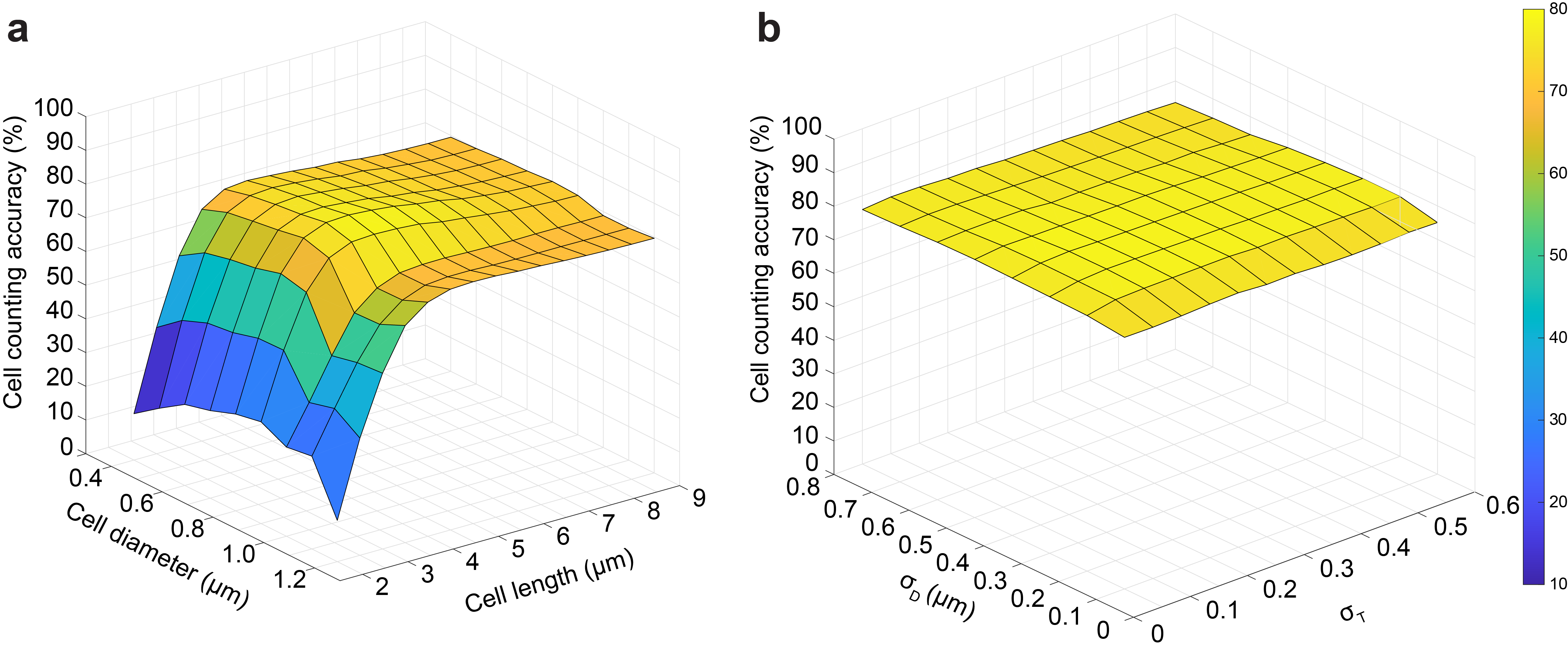

**Figure S13.** **Validation of parameter selection for *LCuts* postprocessing by grid search.** Shown is the cell counting accuracy averaged over 20 randomly chosen, simulated datasets of low SBR and/or high cell density, for which post-processing is required. (**a**) Average cell counting accuracy as a function of cell diameter *d* ∈ [0.4, 1.2] µm and cell length *L* ∈ [2, 9] µm at a fixed *σ_D_* = 0.5 µm and *σ_T_* = 0.2. (**b**) Average cell counting accuracy as a function of *σ_D_* ∈ [0.1, 0.8] µm, and *σ_T_* ∈ [0.05, 0.6] with fixed (*d*, *L*)=(0.8, 4.5) µm. The cell counting accuracy is largely unaffected by variations in *d*, *L*, *σ_D_* and *σ_T_* and robustly remains above 70% for biologically reasonable parameter values, such as *d* ~ 0.8 µm, cell length *L* ~ 6 µm, for *E.coli*-like cell shapes. We also choose *σ_D_* = *d*/2 and *σ_T_* = 0.2, so that edges between nodes separated by more than a cells radius or with relative angles >30° are weighted down.

**Figure S14**

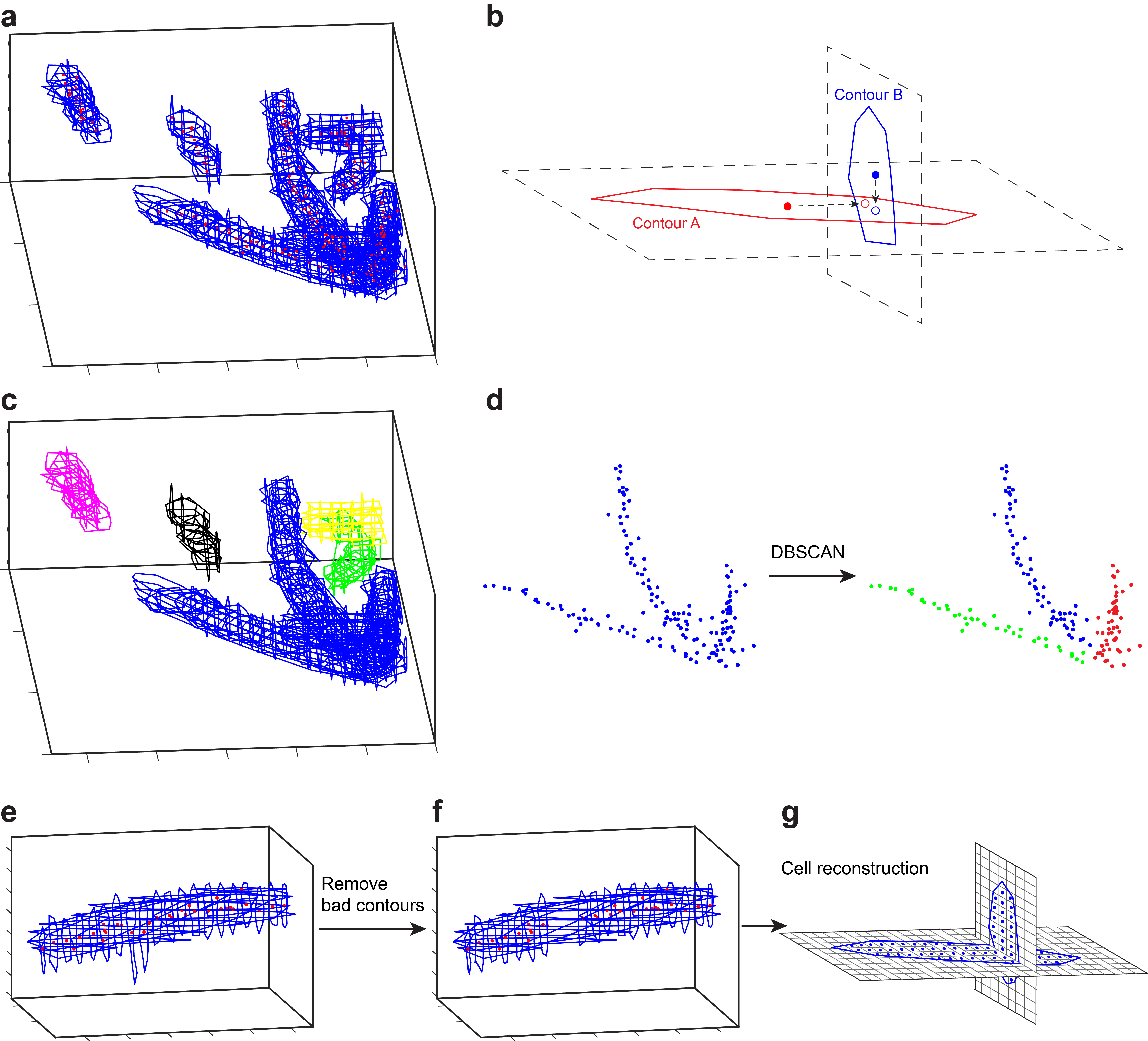

**Figure S14.** **Manual 3D cell shape estimation.** (**a**) Cell shapes in all available x, y and z slices were traced manually. (**b**) Find the contours belonging to the same cell. The centroid of each contour was projected along the x, y and z dimension. If the projected centroid was enclosed by any other contour in a different slice, then the centroid of that contour was projected onto the plane of the initial contour. Two contours were labeled as related if they contained each other’s projected centroids. This process is repeated for all possible contour pairs and their relationship is recorded in an adjacency matrix. (**c**) The related contours are grouped as cells. Different colors represent different cells. (**d**) Segment big clusters that contain more than one cell by grouping the centroids of the contours. This step will run manually and iteratively to segment all single cells from the big cluster. (**e**) Manually check all contours for each cell. (**f**) Remove the bad contours, such as unreasonably large ones. (**g**) A convex hull is built based on the contours for each cell. The convex hull is then used as the mask to extract cell volume from the raw data.
